# Functional Localization of an Attenuating Filter within Cortex for a Selective Detection Task in Mice

**DOI:** 10.1101/2020.03.23.004028

**Authors:** Krithiga Aruljothi, Krista Marrero, Zhaoran Zhang, Behzad Zareian, Edward Zagha

## Abstract

An essential feature of goal-directed behavior is the ability to selectively respond to the diverse stimuli in one’s environment. However, the neural mechanisms that enable us to respond to target stimuli while ignoring distractor stimuli are poorly understood. To study this sensory selection process, we trained male and female mice in a selective detection task in which mice learn to respond to rapid stimuli in the target whisker field and ignore identical stimuli in the opposite, distractor whisker field. In expert mice, we used widefield Ca^2+^ imaging to analyze target-related and distractor-related neural responses throughout dorsal cortex. For target stimuli, we observed strong signal activation in primary somatosensory cortex (S1) and frontal cortices, including both the whisker representation of primary motor cortex (wMC) and anterior lateral motor cortex (ALM). For distractor stimuli, we observe strong signal activation in S1, with minimal propagation to frontal cortex. Our data support only modest subcortical filtering, with robust, step-like attenuation in distractor processing between mono-synaptically coupled regions of S1 and wMC. This study establishes a highly robust model system for studying the neural mechanisms of sensory selection and places important constraints on its implementation.

**Summary:** Responding to task-relevant stimuli while ignoring task-irrelevant stimuli is critical for goal-directed behavior. Yet, the neural mechanisms involved in this selection process are poorly understood. We trained mice in a detection task with both target and distractor stimuli. During expert performance, we measured neural activity throughout cortex using widefield imaging. We observed responses to target stimuli in multiple sensory and motor cortical regions. In contrast, responses to distractor stimuli were abruptly suppressed beyond sensory cortex. Our findings localize the sites of attenuation when successfully ignoring a distractor stimulus, and provide essential foundations for further revealing the neural mechanism of sensory selection and distractor suppression.

## Introduction

We are constantly bombarded by sensory stimuli. To complete a given task, we must selectively respond to task-relevant stimuli while ignoring task-irrelevant stimuli. A framework for understanding stimulus selection is provided by the Treisman attenuation theory (Figure 1). According to this theory, both attended and unattended signals enter short-term storage. Responses to attended stimuli propagate forward for higher-order processing. Responses to unattended stimuli, however, are suppressed by an attenuating filter at some point along the processing stream (Treisman, 1964). The attenuation theory was originally developed to understand selection amongst conflicting speech patterns, yet has since been adapted to study sensory selection across multiple sensory modalities and species (Moran and Desimone, 1985; Wiederman and O’Carroll, 2013; Sridharan et al., 2014).

**Figure 1:**
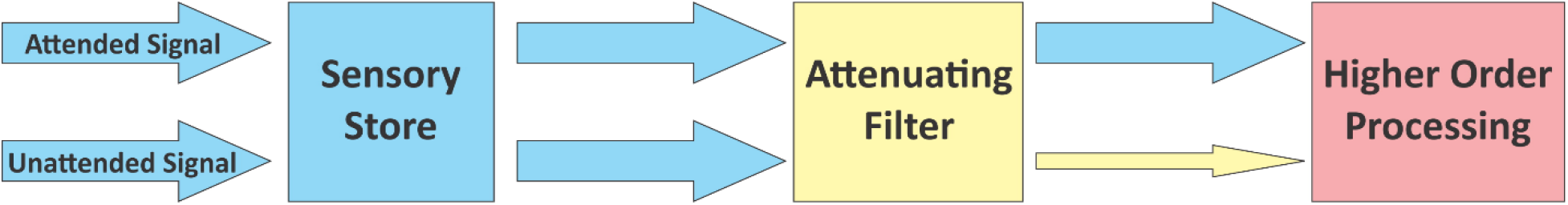
Treisman Attenuation Model. This model of selective attention proposes that both attended and unattended signals enter an early sensory store. At some point in the processing stream, however, an attenuating filter suppresses unattended signals while allowing attended signals to propagate forward for higher order processing.

Where in the brain does attenuation occur and what are the neural mechanisms involved? Extensive studies in the primate visual system have identified stimulus filtering throughout multiple brain regions. Sensory selection was initially proposed to occur in the thalamus, mediated by the modulation of thalamic relay neuron activation by the reticular thalamus (Crick, 1984). Recordings in behaving primates have demonstrated early-onset attentional modulations in thalamus (McAlonan et al., 2008), consistent with stimulus filtering prior to reaching cortex. However, earlier physiological studies demonstrated robust attentional filtering within cortex, between primary visual cortex and visual area V4 (Moran and Desimone, 1985). Alternatively, other studies argue for filtering occurring primarily within prefrontal cortex (Mante et al., 2013). Potential ‘top-down’ pathways establishing an attenuating filter include cortical feedback and ascending neuromodulation (Miller and Cohen, 2001; Noudoost and Moore, 2011). Yet, these mechanisms are poorly understood, in part due to the apparent highly distributed filtering processes of the primate visual system.

Our goal in this study is to localize the attenuating filter for a simple sensory-motor task in the mouse whisker system. This model system benefits from extensive characterization of the sensory and motor regions and pathways involved, with significantly fewer hierarchical levels than the primate visual system (Kleinfeld et al., 1999; Guo et al., 2014; Petersen, 2019). Whisker deflection activates brainstem pathways which travel predominantly through the ventral posteromedial (VPM) thalamus and onto primary somatosensory (barrel) cortex (S1). From S1, there are robust, mono-synaptic connections to the whisker representation of primary motor cortex (wMC) (Porter and White, 1983; Miyashita et al., 1994; Mao et al., 2011). Sensory responses in S1 rapidly propagate to wMC, under both anesthetized and awake conditions (Farkas et al., 1999; Kleinfeld et al., 2002; Ferezou et al., 2007; Chakrabarti et al., 2008; Zagha et al., 2015). Moreover, this pathway may be particularly important for input detection; S1-wMC projection neurons were found to be preferentially responsive to touch in an object detection task (Chen et al., 2013), which enhanced during task training (Chen et al., 2015). Recent studies in the rodent whisker system have reported sensory filtering within the thalamus (Rodenkirch et al., 2019) and brainstem (Chakrabarti and Schwarz, 2018). However, it remains unknown to what extent these subcortical or cortical pathways contribute to filtering during a sensory selection task.

We designed a selective detection task with spatially and temporally distinct processing streams. Mice respond to rapid deflections of one whisker field (target) and ignore identical stimuli in the opposite, contralateral whisker field (distractor). Rather than presenting target and distractor stimuli together, as in the original studies on sensory selection (Treisman, 1964; Moran and Desimone, 1985), we present each stimulus individually on different trials. Thus, we can evaluate target and distractor processing separately across space (different hemispheres) and time (different trials). The motor response in our task is a straight-forward lick. As the sensory and motor content of our task is symmetric, the only asymmetry is the selection process. In expert performing mice, we used widefield Ca^2+^ population imaging (Wekselblatt et al., 2016) to simultaneously monitor neural activity bilaterally in sensory and motor regions. We then quantified the asymmetry in target-aligned versus distractor-aligned sensory processing streams to localize sites of attenuation.

## Materials and Methods

### Animal Subjects and Surgery

All experiments performed in this study were approved by the IACUC of University of California, Riverside. Mice were purchased from Jackson Laboratories (JAX). Task-related neural imaging data were obtained from GCaMP6s expressing Snap25-2A-GCaMP6s-D mice (JAX #025111). The SNAP25-2A-GCaMP6s mouse line expresses GCaMP6s pan-neuronally, in both excitatory and inhibitory neurons throughout the brain (Madisen et al., 2015). Transgenic mice were backcrossed into the BALB/cByJ (JAX 000651) background. Both male and female mice were used in these experiments. Recording sessions from male and female mice were similar according to behavioral performance (imaging experiments: 4 male mice, 32 sessions, 1 female mouse, 7 sessions; discriminability d’: male 2.0 ± 0.1, female 1.9 ± 0.2, two sample t-test p= 0.36; target stimulus reaction time (s): male 0.30 ± 0.01, female 0.32 ± 0.02, two sample t-test p= 0.70) and neural responses (data not shown), and therefore data were combined for grand average analyses. Mice were housed on a light cycle of 12 hours light/ 12 hours dark. All trainings and recordings were conducted on mice head-fixed in the behavioral apparatus. For headpost implantation, 2 to 5 months-old mice were placed under a combination of isoflurane (1-2%), ketamine (100 mg/kg), and xylazine (10 mg/kg) anesthesia. A 10 mm x 10 mm piece of scalp was resected to expose the skull. The exposed skull was cleared of connective tissue and a custom-built headpost was implanted onto the skull with cyanoacrylate glue. The lightweight titanium or stainless steel headpost (3 cm in length and 1.5 grams in weight) had an 8 mm x 8 mm central window for imaging and recording. For in vivo widefield Ca^2+^ imaging, a thin layer of cyanoacrylate gap-filling medium (Insta-Cure, Bob Smith Industries) was applied to the window, to both seal the exposed skull and enhance skull transparency. Silicone elastomer (Reynolds Advanced Materials) was additionally applied above the imaging window. After surgery, mice were placed onto a heating pad to recover and administered meloxicam (0.3 mg/kg) and enrofloxacin (5 mg/kg) for three days post-op. Mice were given a minimum of three days to recover from surgery before water-restriction and behavioral training. Recordings under anesthesia were conducted immediately after headpost implantation.

### Animal Behavior

Mice were trained in a Go/NoGo passive whisker selective detection task. During behavioral training mice were given food ad libitum but were water-restricted to a minimum of 1 mL per day. Weights were monitored daily to maintain over 85% of their initial post-surgery weights, and additional water was given as needed to maintain this level. The behavioral apparatus was controlled by Arduino and custom MATLAB (MathWorks) code. Piezo-controlled paddles (Physik Instrumente and Piezo.com) were placed bilaterally in the whisker fields, with each paddle contacting 2 to 4 whiskers. Paddle deflections of a triangle waveform had rising phases that ranged from 0.1 s (for large deflections) to 0.01 s (for small deflections), followed by an immediate falling phase. Deflection velocity was constant, therefore increased duration correlated with increased deflection amplitude. The maximum amplitude, for 0.1 s deflections, was 1 mm. Stimulus duration and amplitude were varied with training with the goal of maintaining a 75% hit rate. This target hit rate was selected to maintain high reward rates while still operating within the dynamic range of each mouse’s psychometric curve. Within every session, target and distractor stimulus strengths were identical. Directly below the mouse’s snout was a central lick port. Each ‘hit’ trial was rewarded with ~5 μL of water delivered through the lick port.

Behavioral training consisted of three stages. Inter-trial intervals for all stages varied from 5 to 9 s with a negative exponential distribution to minimize timing strategies. Additionally, in all stages a ‘lockout’ period of 200 ms separated stimulus onset and the earliest opportunity for reward. Target and distractor whisker fields were assigned at Stage 1 and remained constant throughout training. Target/distractor assignment was varied across the population and analyzed separately (Figure 7A-C) before combining for grand average analyses. Each session lasted approximately 60 minutes and consisted of 100 to 200 trials. (Stage 1) Classical conditioning: Unilateral (target) whisker deflection was paired with fluid reward; distractor whisker deflection was neither rewarded nor punished. Mice were trained on this stage for 2 to 3 days, 1 to 2 sessions per day. (Stage 2) Operant conditioning: Following unilateral (target) whisker deflection, mice were required to contact the lick port within a lick detection window of 1.5 s in order to initiate the fluid reward. Mice were trained on this stage for 2 to 3 days, 1 session per day. (Stage 3) Impulse control: Similar task structure as above, except all incorrect responses (licking during the ITI, during the lockout period, or following distractor deflections) were punished by re-setting the ITI, effectively acting as a time-out. The response detection window was shortened to 1 s. Following full-length ITIs, trial types were selected randomly from a distribution of 80% distractor and 20% target. For distractor trials, not responding *(correct rejection)* was rewarded with a shortened ITI (2 to 4 s, negative exponential distribution) and a subsequent target trial. Licking to the distractor *(false alarm)* or not responding to the target *(miss)* initiated a subsequent full-length ITI. Responding to the target stimulus *(hit)* triggered a fluid reward, followed by a full-length ITI. Behavioral and neural imaging data for hit trials with and without preceding correct rejections were compared (Figure 7D, E) before combining for grand average analyses.

A single, contiguous behavioral window was considered for analyses, from session onset until 120 s of no responding, which we interpreted as task disengagement. Hit rate, false alarm rate, spontaneous lick rate, and reaction times were all used to assess task performance. Foremost, we used the sensitivity or d-prime (d’) framework from signal detection theory. Traditionally, d’ is used as a measure of detection between stimulus present and stimulus absent conditions. Here, we implemented a discriminability d’ between target detection and distractor detection [d’ = Z_hit rate_ – Z_false alarm rate_] where Z is the inverse of the normal cumulative distribution function. Mice were considered expert in our task once they achieved a d’>1 for three consecutive days. Spontaneous lick rate was calculated as the response rate during the last 1 s of the full-length ITI.

### Widefield Imaging

Widefield imaging was performed through-skull in head-fixed mice while they performed the selective detection task. Imaging was conducted through a Macroscope IIa (RedShirtImaging), beam diverter removed, 75 mm inverted lens with 0.7x magnification and 16 mm working distance. The lens was positioned directly over the cranial window, providing a 7 mm x 5.8 mm field of view, including most of dorsal parietal and frontal cortex bilaterally. Illumination was provided by a mounted 470 nm LED (Thorlabs M470L3), dispersed with a collimating lens (Thorlabs ACL2520-A), band-pass filtered (Chroma ET480/40x) and directed through the macroscope using a dichroic mirror (Chroma T510lpxrxt). Fluorescent light returning to the brain was band-pass filtered (Chroma ET535/50m) prior to reaching an RT sCMOS camera (SPOT Imaging). On camera 2×2 binning and post-processing image size reduction gave a final resolution of 142 x 170 pixels at 41 μm per pixel and 12-bit depth. Images were acquired at a temporal resolution of 10 Hz, aligned to the trial structure. TIF image sequences were imported to MATLAB for preprocessing and analysis.

### Local field potential (LFP) recordings

LFP recordings were conducted through small (<0.5 mm diameter) craniotomies and duratomies positioned above S1 (from bregma: posterior 1.5 mm, lateral 3.5 mm), wMC (anterior 1 mm, lateral 1 mm) and ALM (anterior 2.5 mm, lateral 1.5 mm), in target-aligned and distractor-aligned cortices. Recording sites were positioned 900 μm below the pial surface, targeting layer 5. Recordings were acquired with silicon probes (Neuronexus, A1×16-Poly2-5mm-50s-177), bandpass filtered from 0.1 Hz to 8 kHz and digitized at 32 kHz (Neuralynx). Further analyses were conducted in MATLAB.

### Imaging of whisker movements

A CMOS camera (Thorlabs DCC3240M camera with Edmund Optics lens 33-301) was positioned directly above the mouse while performing the detection task. Field of view included both whisker fields and stimulus paddles. Images were captured at 8-bit depth continuously at 60 Hz (ThorCam) and imported to MATLAB for analyses.

### Data Analysis

All data analyses were performed in MATLAB using custom scripts.

### Fluorescence Preprocessing and Trial-Based Neural Activity

Peri-stimulus trial imaging time windows included 1 s before stimulus onset and 1.2 s after stimulus onset, which included the lockout and response windows. The first step of image processing was to concatenate fluorescence activity from consecutive trials to create a raw movie F, where F_n_(i,j,f) shows the fluorescence of each pixel (i^th^ row, j^th^ column) in the f^th^ frame for each individual trial n. The pre-stimulus baseline fluorescence F_o_(i,j,n) was calculated by averaging pixelwise activity across the first 10 frames preceding the stimulus onset per trial n (1 s pre-stimulus). Finally, relative fluorescent signal normalized to pre-stimulus baseline (dF/F) was calculated as

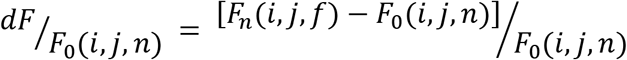

Average trial movies were created by indexing trials according to trial outcome (*hit, miss, false alarm, correct rejection, spontaneous*) and averaging activities at each pixel across the corresponding frame of each corresponding trial. Frame alignments were conducted both with reference to stimulus onset (stimulus-aligned) and with reference to the first frame containing the response (response-aligned) (see Figure 4). Spontaneous trials were those in which a response occurred during the 1 s pre-stimulus imaging period. Trials with responses during the lockout period were excluded from all further analyses.

Data were analyzed per session (n=39), per mouse (n=5), per target-distractor assignment (n=2) and across all experiments (grand average). For a session to be included in our analyses, our inclusion criteria were d’>1 for at least 10 minutes of continuous engagement. Only one engagement period per session was included. For qualitative analyses, trial movies from recording sessions were spatially aligned to bregma and averaged per mouse. These data were then averaged per target-distractor assignment (see Figure 7 for whisker deflection assignments). One target-distractor assignment dataset was then flipped horizontally (rostro-caudal axis) at bregma before the grand average dF/F. For quantitative analyses, the subsequent datasets were first flipped at bregma according to target-distractor assignment (as before) and then averaged across all sessions.

### Quantification of Stimulus Encoding

To quantify stimulus response magnitude, we calculated the neurometric d’ (Britten et al., 1992) comparing activity pre-stimulus (stimulus absent) and post-stimulus (stimulus present), specifically during the lockout period. Neurometric d’ was calculated separately for target and distractor trials, and included all trials regardless of outcome (hit and miss trials for target, false alarm and correct rejection for distractor). Pre-stimulus (10 frames preceding stimulus onset) and post-stimulus activities were binned and plotted in an ROC (receiver operating characteristic) curve. The area under the curve (AUC) was converted to d’ using the equation:

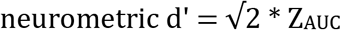

In the context of this study, neurometric d’ is the performance measurement of a pixel where d’>0 denotes more post-stimulus pixel activity and d’<0 denotes more pre-stimulus pixel activity. For each region of interest, we report the peak neurometric d’ within the spatially-defined region of interest (ROI). Subsequent analyses compared target stimulus encoding in target-aligned cortices to distractor stimulus encoding in distractor-aligned cortices.

### Quantification of Choice Probability

To quantify choice-related neural activity, we calculated choice probability d’ (Britten et al., 1996) comparing activity on hit trials (response present) and miss trials (response absent), specifically during the lockout phase. Sessions were included in this analysis if they had 5 or more trials of each type. Trials with stimuli of different amplitudes were combined only if response rates for each amplitude-specific trial type were comparable (within 15%). Overall, 9 sessions were excluded from this analysis, due to too few miss trials (n=30, instead of n=39). Choice probability was calculated for activity within the pre-response frame (100 to 200 ms during the lockout) and for activity between the pre-response frames (change in activity, subtraction of activity in the 0 to 100 ms frame window from the 100 to 200 ms frame window during the lockout). In the context of this study, choice probability d’ is the performance measurement of a pixel where d’>0 denotes more response-related pixel activity and d’<0 denotes more no response-related pixel activity. Response present and response absent activities were binned and plotted in a ROC curve. The area under the curve was converted to d’ as described above.

### Seed Correlation Analysis

Correlation maps were generated separately for target and distractor hemispheres and for S1, wMC and ALM seed regions (generating six correlation maps per session). Baseline averaged fluorescence activity trajectories from all trial types (excluding spontaneous) were concatenated into a single time series. The following trial structures were analyzed separately: 1) full trial, including 10 frames pre-stimulus and 12 frames post-stimulus including the lockout and response windows, 2) pre-stimulus only, including 10 frames pre-stimulus, 3) peri-stimulus and lockout, including 1 frame pre-stimulus and 2 frames post-stimulus during the lockout, and 4) response, including 10 frames after the lockout and during the response window. The seed was the average time series from all pixels in the indicated region of interest. Pairwise correlation coefficients were calculated between the seed and all other pixels. To reduce computation time, all trial movies were spatially down sampled 4-fold across both axes for a resolution of 36 x 43 pixels at 160 um per pixel prior to running the correlation analyses. We report r^2^ values, as the square of the correlation coefficient. For each region of interest, we report the average correlation r^2^ within the spatially defined region. Subsequent analyses compared target-aligned intracortical correlations to distractor-aligned intracortical correlations.

### Evoked-potential analyses

Single-trial LFP recordings were aligned to target and distractor stimulus onset. To isolate the LFP signal, single-trial data were bandpass filtered from 0.2 Hz to 100 Hz using a second order Butterworth filter, then downsampled to a sampling frequency of 400 Hz. Following filtering and downsampling, single-trial data were averaged according to trial type. In figure 9, stimulus artifacts at 0-10 ms post-stimulus were truncated when present.

### Whisker movement analyses

Movies were parsed into regions of interest containing target or distractor whisker fields. Whisker motion energy (WME) within each region was calculated for each frame as the temporal derivative for each pixel of the mean gray value from the previous frame. Values per pixel were normalized (squared) and summed across pixels, providing a single WME value. WME data from the movies were aligned to target and distractor stimulus onset and averaged across trial type.

### Statistical Analyses

For neurometric d’ and choice probability d’, statistical analyses were performed to determine whether each pixel value was significantly different than zero across sessions (one sample t-test). Data were spatially aligned across sessions as described above. Threshold for statistical significance was corrected for multiple comparisons using the Bonferroni correction [0.05/(142×170) = 2.1×10^-6^ for a single imaging session and 0.05/(156×194) = 1.7×10^-6^ across aligned imaging sessions]. For neurometric d’ and seed correlation, we additionally conducted region of interest (ROI) analyses. For neurometric d’, reduction in distractor encoding was calculated as: (target d’ – distractor d’) / target d’, calculated separately for S1, wMC and ALM. Statistical analyses were performed to determine whether reduction in distractor encoding was significantly different than zero within each region across sessions (one sample t-test, significance threshold corrected for multiple comparisons 0.05/3 = 0.017). Additionally, comparison of reduction in distractor encoding between the three ROIs across sessions was conducted using ANOVA and post-hoc Tukey test. For seed correlation, comparison between the three ROIs across sessions and comparisons between different trial phases across sessions were conducted using ANOVA and post-hoc Tukey test. To quantify changes in whisker motion energy (WME), post-stimulus values (each frame) were compared to average prestimulus (1 s baseline) values. Comparisons were conducted using paired t-test for each post-stimulus window, with a p-value threshold of 0.01 for significance. Average data are reported as mean ± standard error of the mean.

## Results

### Training mice in a selective detection task

To study the neural mechanisms of sensory selection, we developed a Go/NoGo passive whisker detection task in head-fixed mice (Figure 2). In this task, target stimuli are rapid deflections of multiple whiskers in one whisker field and distractor stimuli are identical deflections in the opposite whisker field (Figure 2A). Throughout training we quantify task performance as the separation between hit rate and false alarm rate (Figure 2C). We considered mice ‘expert’ once they achieved a discriminability d’>1 on three consecutive sessions. Average time to expert performance was 11 days in the full task (see Methods) (Figures 2D and 2E) (number of sessions to expert performance: 11.2±0.9, n=43 mice). Performance measures for the imaging sessions used in subsequent analyses are shown in Figure 2F (n=39 sessions across n=5 mice, hit rate (%), 80.4 ± 2.2; false alarm rate, 13.6 ± 1.1; spontaneous lick rate, 8.1 ± 0.5; d’ comparing hit vs false alarm rates, 2.0 ± 0.1).

**Figure 2:**
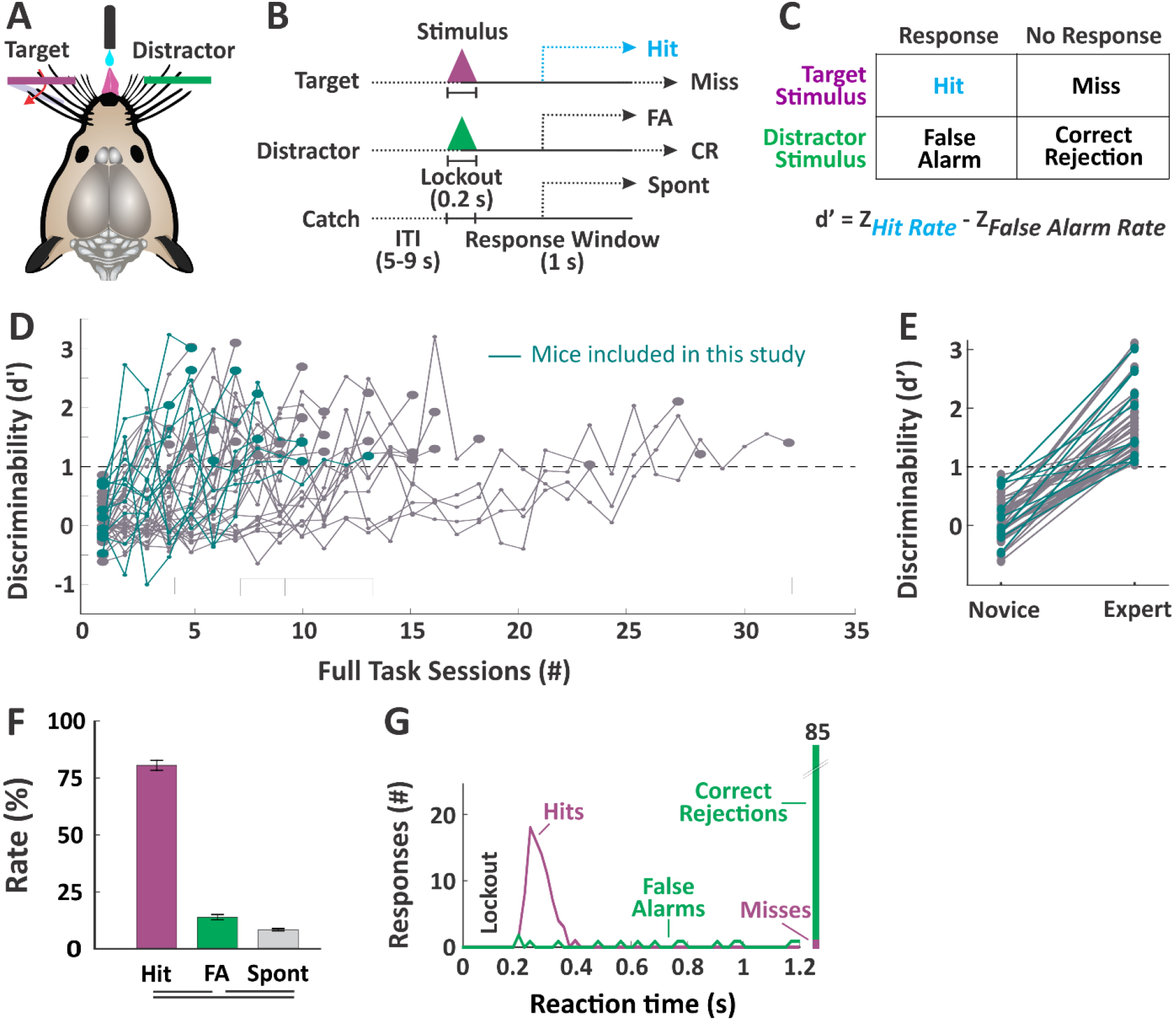
Behavior Paradigm and Measures of Selective Detection. **(A)** Illustration of the behavioral setup. Mice are head-fixed into the behavioral rig with piezo-controlled paddles within their whisker fields bilaterally. Each paddle is assigned as target (purple) or distractor (green) at the start of training. Mice report stimulus detection and receive rewards from a central lickport. **(B)** Task structure. Each trial consists of an inter-trial interval, a stimulus and 200ms lockout, and a 1 s response window. Trial type as determined by the stimulus could be target, distractor or catch (no stimulus). **(C)** Calculation of discriminability d’, as the separation between hit rate and false alarm rate. **(D)** Performance trajectories for all mice (n=43 mice) and box and whiskers summary plot. Those used for imaging studies (n=5 mice) are indicated in blue. Mice were considered expert once they achieved a d’>1 for three consecutive days. **(E)** Comparison of d’ for novice mice (first day of training on impulse control) and expert mice (n=43 mice, p<10^-19^). **(F)** Performance measures for the imaging sessions (n=39 sessions). Lines below plot denote statistical significance. **(G)** Example session data showing reaction time distributions for target and distractor trials.

Two key features of this task facilitate the study of sensory selection. First, target and distractor stimuli are presented to contralateral whisker fields. Given the highly lateralized somatosensory whisker representation, we expect the target-aligned and distractor-aligned processing streams to be well separated across hemispheres. Second, we imposed a short (200 ms) lockout period after stimulus onset and before the response window. Responding during the lockout is punished with a time-out, and mice learn to withhold their responses through this period (e.g., Figure 2G). All analyses of stimulus selection are conducted within this lockout period, which is post-stimulus onset and pre-response, thereby isolating the selection process from overt behavior.

### Propagation of cortical activity during task performance

We used widefield calcium imaging (GCaMP6s Ca^2+^ sensor) to monitor neural activity broadly across dorsal cortex during task performance. We used a combination of anatomic landmarks and functional mapping to identify various cortical regions (Figures 3A and 3B). Whisker deflection in anesthetized mice was used to localize the primary somatosensory barrel field (S1) and the whisker representation of primary motor cortex (wMC) (n=13, example session shown in Figure 3B, left). Reward-triggered licking in water-restricted yet task naïve mice was used to localize anterior lateral motor cortex (ALM), which has recently been identified as a pre-motor licking-related region (Guo et al., 2014; Chen et al., 2017) (n=6, example session shown in Figure 3B, right). Thus, our anatomic and functional mapping confirm that we can simultaneously monitor licking-related and whisker sensory and motor cortical regions bilaterally.

**Figure 3:**
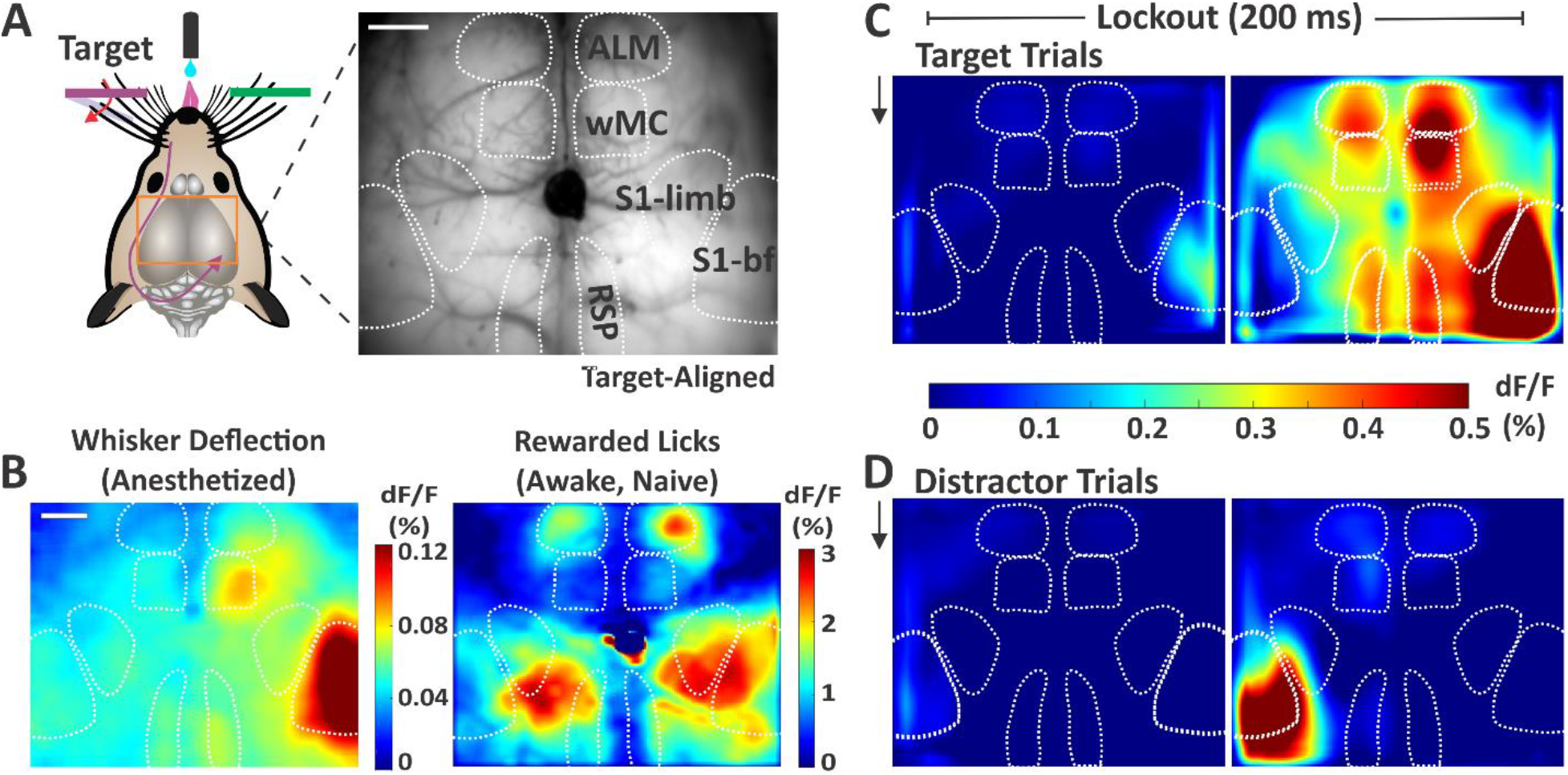
Sensory and Motor Cortical Representations Using Widefield Ca^2+^ Imaging. **(A)** Illustration of the imaging setup (left) and example frame from the through-skull GCaMP6s imaging (right). Surface vessels appear as dark striations overlaying the brain parenchyma. Bregma is indicated by the central ink blot. **(B)** Cortical activity (dF/F) following whisker deflections in an anesthetized mouse (left), to localize of the sensory and motor whisker representations. Cortical activity following reward-triggered licking in a naïve mouse (right), to localize licking-related activity. **(C)** Cortical activity on target trials during the two sequential imaging frames of the lockout period in expert mice performing the detection task (grand average, n=39 sessions). Black arrow indicates whisker stimulus onset, which is coincident with the start of the first imaging frame. **(D)** Same as [C], but for distractor trials. Note the differential propagation of cortical activity depending on trial type. Scale bars in [A] and [B] are 1mm.

We imaged expert mice while they were performing the whisker detection task. Here, we show stimulus evoked cortical activity on target and distractor trials across all mice and all sessions (grand average: n=5 mice, n=39 sessions) (Figures 3C and 3D). The two sequential imaging frames both occur within the lockout period, which is after stimulus onset and before the earliest allowed response time. As expected, for both trial types we observe activity initiation in S1 contralateral to the deflected whisker field. By the end of the lockout period we observe strong S1 activity following both target and distractor stimuli. On target trials we observe propagation of activity to wMC, ALM, and retrosplenial cortex (RSP). Note that the activity does not spread uniformly from the site of initiation, but rather emerges in discrete cortical regions. In contrast, on distractor trials the activity is largely contained within S1, with only mild activation of wMC.

In Figure 4, we show the grand average fluorescence signals across all trial types and outcomes, aligned to both stimulus onset and response onset. Notice that during the response (post-response onset for hit, false alarm and spontaneous licking trials) we see strong signals that are widespread throughout dorsal cortex. However, in this study we are most interested in the activity initiating, and therefore preceding, the response. On hit trials (Figure 4A), we observe the propagation of activity from S1 to frontal and parietal regions post-stimulus (aligned to stimulus) and pre-response (aligned to response). On correct rejection trials (Figure 4E), we also see strong activity in S1, but with very little propagation to other cortical regions. Propagation is not simply delayed on these trials, as we can track the resolution of distractor-evoked activity into the response window.

**Figure 4:**
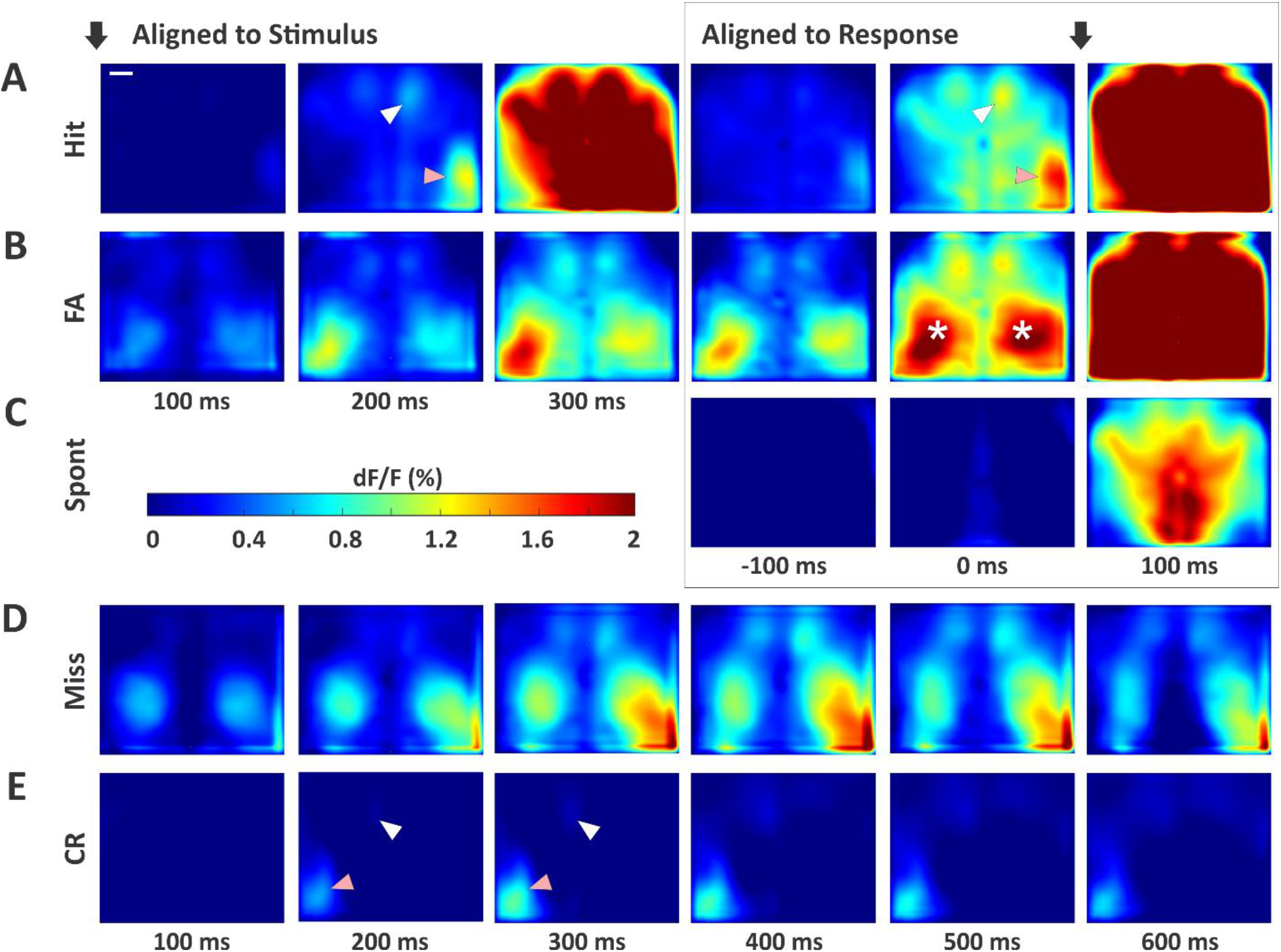
Cortical Activity Patterns Across All Trial Types. **(A)** Hit trials. Black arrows indicate alignment to stimulus onset (left three panels) or response onset (right three panels). The third frame aligned to stimulus (300 ms) is the first frame after the lockout and within the response window. Note the strong activity in contralateral S1 (pink arrows) with propagation to wMC (white arrows) and ALM, prior to response generation. **(B)** False alarm trials, with the same plot structure as in [A]. Asterisks mark elevated activity in the S1-limb regions, bilaterally. **(C)** Spontaneous trials (no stimulus alignment). **(D)** Miss trials. As there is no response on these trial types, we plot an extended series of post-stimulus activity. **(E)** Correct rejection trials, with the same plot structure as in [D]. Note the strong activity in S1 (pink arrow), yet lack of propagation to wMC (white arrow) and ALM. Scale bar in [A] is 1 mm.

The incorrect trial types also show distinct activation patterns. On both false alarm trials (Figure 4B) and miss trials (Figure 4D), in addition to lateralized S1 responses, we also observe prominent bilateral activity in the somatosensory limb regions. We interpret these neural signals as reflecting self-motion of the mouse. Prior studies have shown that during passive whisker detection tasks, self-motion (quantified by whisking behavior) reduces detection probability (Ollerenshaw et al., 2012). Thus, limb region activation observed here is consistent with self-motion contributing to incorrect, both miss and false alarm, trial outcomes. On spontaneous trials (responses not preceded by a whisker stimulus) we observe minimal pre-response cortical activity (Figure 4C).

### Quantification of stimulus encoding and attenuation across cortex

The above analyses demonstrate, qualitatively, the differential propagation of cortical signals for target and distractor stimuli. Next, we sought to quantify these responses. To do this, we calculated the neurometric sensitivity index (d’) (Britten et al., 1992) for each pixel in our imaging window (Figure 5). Across each session we compared the prestimulus activity (stimulus absent) to activity during the lockout period (stimulus present). Importantly, for this analysis we included all target trials and all distractor trials regardless of trial outcome. We use d’ rather than dF/F, as the former accounts for trial-by-trial variability and reflects the ability of an ideal observer to distinguish signal from noise on single trials. The d’ maps from target and distractor stimuli largely match dF/F patterns described above; for target stimuli high d’ values are observed in S1, wMC, ALM and RSP (Figure 5A, right) whereas for distractor stimuli, high d’ values are only observed in S1 (Figure 5B, right). These regions show neurometric d’ values significantly above zero (Figure 5C and 5D). We do observe a focal increase in d’ for distractor wMC (Figure 5B, right), but this does not reach statistical significance after correction for multiple comparisons (Figure 5D).

**Figure 5:**
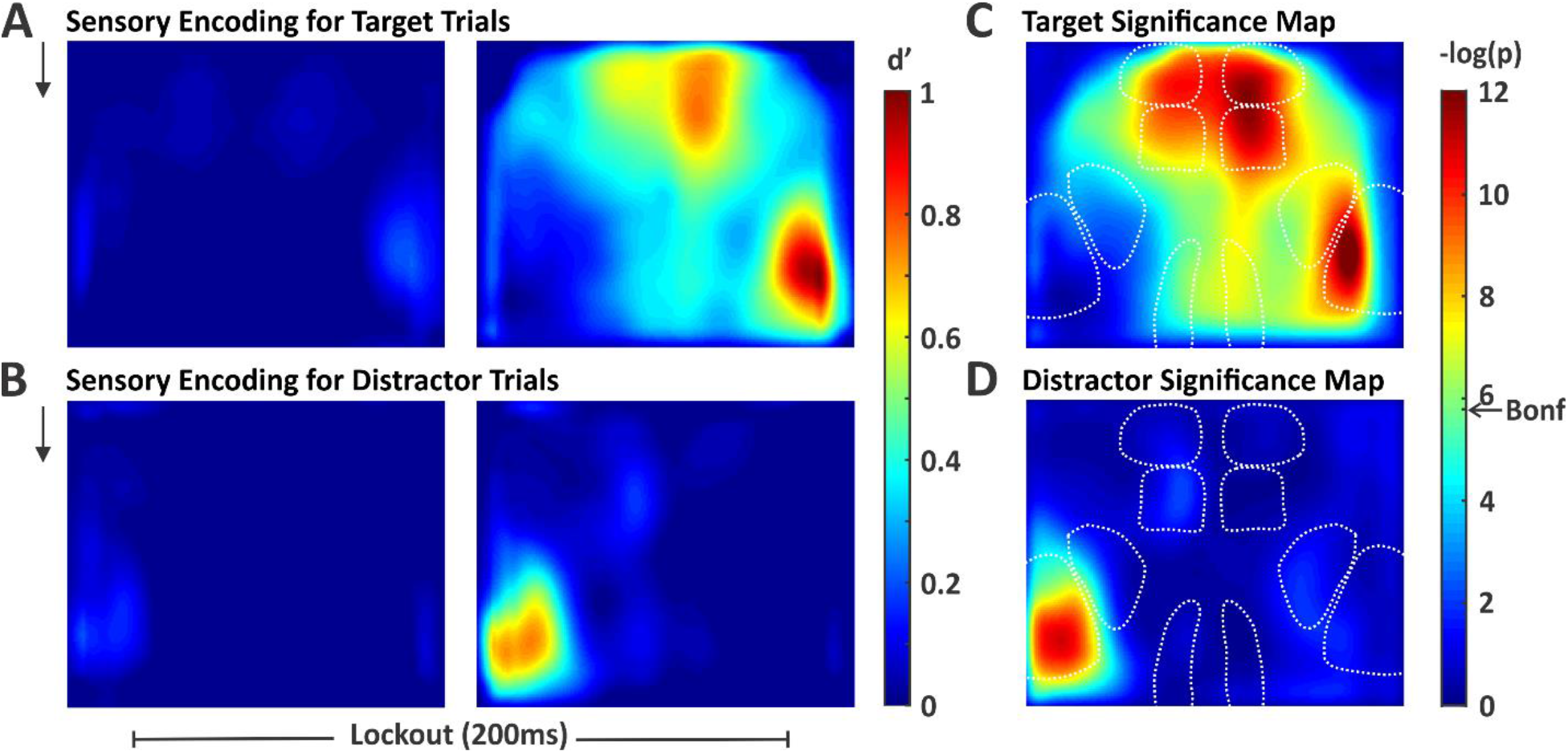
Spatial Maps of Stimulus Encoding. We quantified stimulus encoding as the separation between stimulus absent and stimulus present d’, computed pixel-by-pixel. **(A)** Map of target stimulus encoding during the two sequential frames of the lockout period (black arrow represents stimulus onset). **(B)** Map of distractor stimulus encoding during the same time windows as in [A]. **(C)** and **(D)** Significance maps of the right panels of [A] and [B], respectively. Significance threshold determined by the Bonferroni correction for multiple comparisons is indicated by the arrow on the color bar (Bonf). Pixels with smaller p-values (warmer colors) have d’ values significantly above 0. For target stimuli, we observe widespread stimulus encoding including in multiple frontal and parietal regions. For distractor stimuli, significant stimulus encoding is restricted to S1.

Next, we quantify the propagation of stimulus responses for target versus distractor stimuli. We describe this analysis first for S1. For each session, we determined the peak neurometric d’ for the target stimulus in target-aligned S1 versus the peak neurometric d’ for the distractor stimulus in distractor-aligned S1. We plot these data in Figure 6A. Data along the unity line indicate equal neurometric d’ values for target and distractor stimuli for that session. For S1, the data are widely distributed, yet with a nonsignificant trend towards larger responses for target stimuli (n=39 sessions, 8.8 ± 7.9% reduction in distractor d’, p=0.27 (t(38)= 1.12, one sample t-test) (Figure 6D). We repeated these analyses for wMC and ALM. For these regions we find that neurometric d’ values are consistently larger for target stimuli (Figures 6B and 6C). Reduction in distractor d’ is 61.0±7.1% in wMC (p=1.76e^-10^, t(38)= 8.62, one sample t-test) and 72.1 ± 6.9 % in ALM (p=8.83e^-13^, t(38)= 10.49, one sample t-test) (Figure 6D). Additionally, reduction in distractor encoding is greater for wMC and ALM compared to S1 (p=7.97e^-9^, F(1,38) = 54.2, ANOVA with post-hoc Tukey comparison). Overall, these data demonstrate robust attenuation of distractor responses between the mono-synaptically connected regions of S1 and wMC.

**Figure 6:**
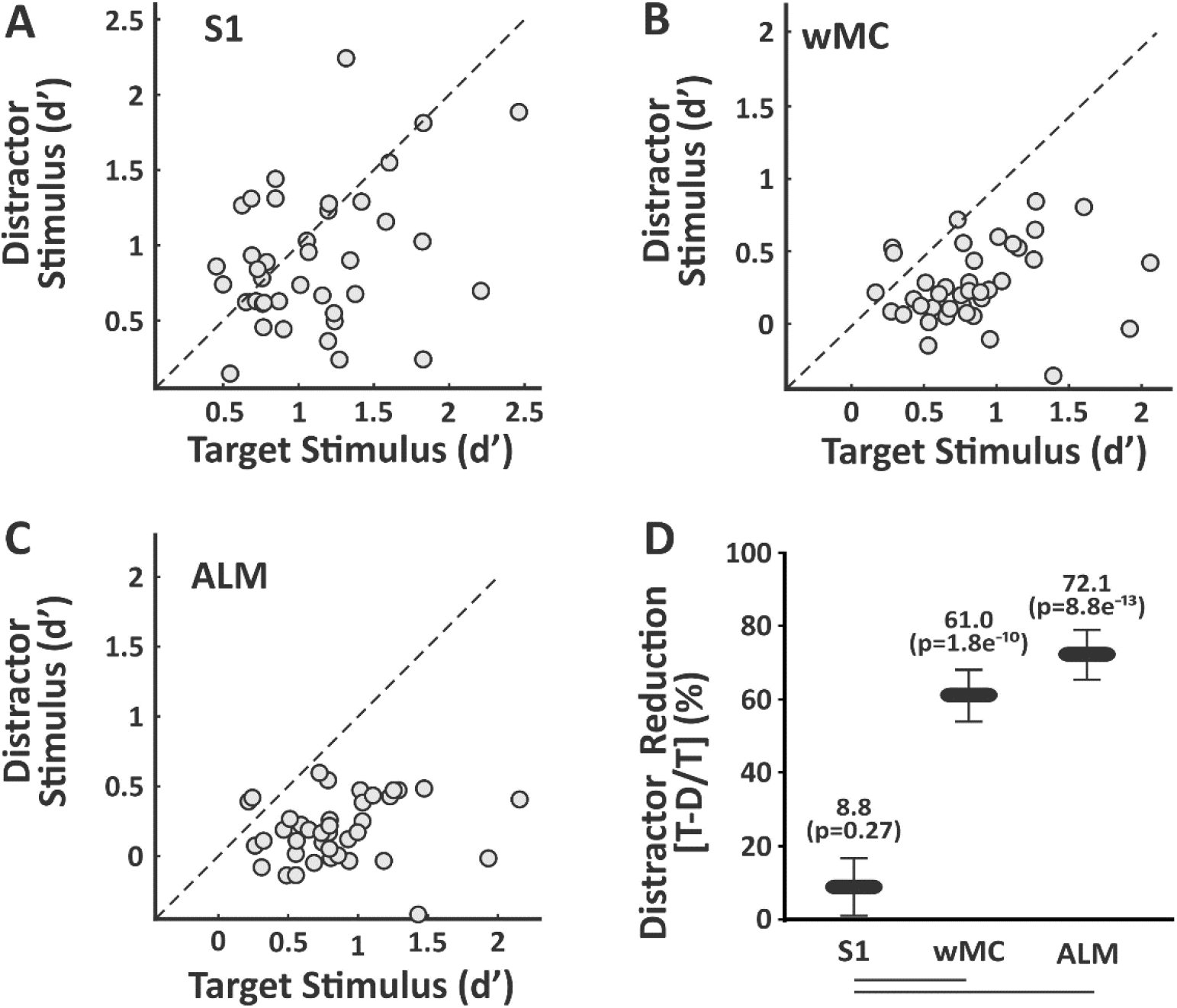
Quantification of Target vs Distractor Stimulus Propagation within Cortex. For each session, we compared target stimulus encoding in target-aligned cortices to distractor stimulus encoding in distractor-aligned cortex. **(A-C)** Scatter plots of target versus distractor encoding in S1 **(A)**, wMC **(B)** and ALM **(C)**. Each data point is one session (n=39 sesssions). Note that the data are broadly distributed in S1, and highly biased towards stronger target encoding in wMC and ALM. **(D)** Summary data, comparing reductions in distractor encoding within each region (values above each data point) and between regions (lines below graph denote statistical significance). Reductions in distractor encoding are significantly larger in wMC and ALM compared to S1.

### Analyses of intrinsic lateralization, trial history, electrical activity, whisker movements and choice probability

We performed a series of analyses to determine whether the neural activity described above reflects the selection process or can be accounted for by task or behavioral confounds. First, widespread cortical propagation in our task could reflect target selection or an intrinsic lateralization of cortical activity (e.g., left-sided whisker deflections always evoke more widespread cortical activation). To distinguish between these possibilities, two cohorts of mice were trained with opposite target-distractor assignments. In previous analyses we aligned all data with respect to target-distractor orientation. Here, we show behavioral performance (Figure 7A) and neural activity separately according to target assignment (Figure 7B target-aligned right hemisphere, n=3 mice and n=25 sessions; Figure 7C target-aligned left hemisphere, n=2 mice, n=14 sessions). Note that propagation from S1 to frontal and other parietal cortices occurs selectively on target trials, irrespective of the side of target assignment. Therefore, these differential patterns of cortical activation reflect learned adaptations to our task, rather than intrinsic lateralization.

**Figure 7:**
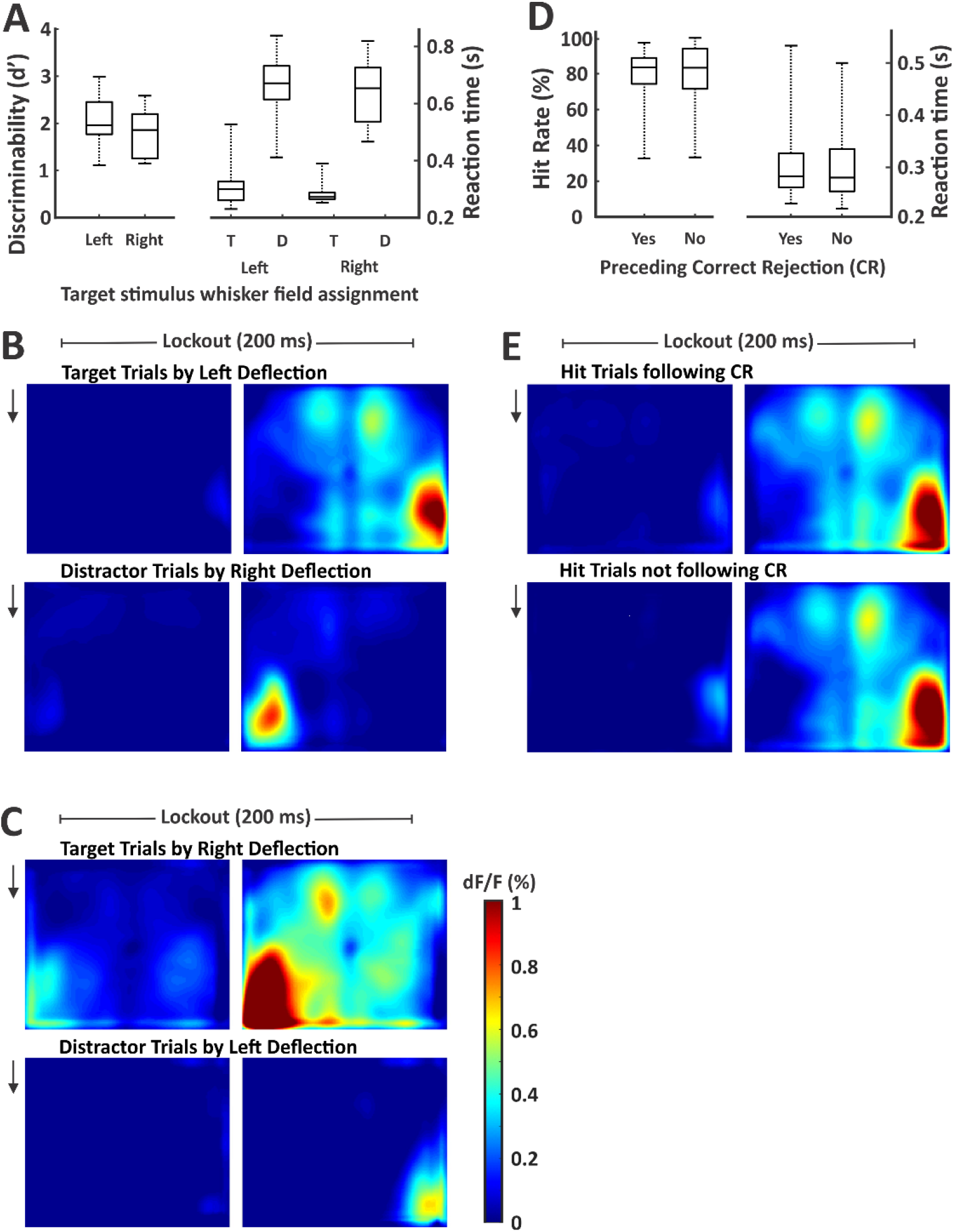
Similar behavior and neural activity across target assignments and trial structures. **(A)** Discriminability d’ and reaction times reported separately for mice with left or right target whisker field assignment. None of the behavioral measures were significantly different between these two populations. **(B)** Cortical activity during the lockout period for target trials (top) and distractor trials (bottom) for sessions in which the target was assigned to the left whisker field (represented by the right cortical hemisphere) (n = 3 mice, n = 25 sessions). **(C)** Same as [B], but for sessions in which the target was assigned to the right whisker field (represented by the left cortical hemisphere) (n = 2 mice, n = 14 sessions). Signal propagation to frontal cortex correlated with target assignment. **(D)** Hit rates and reaction times reported separately for target trials with and without a preceding correct rejection. None of the behavioral measures were significantly different between these two trial structures. **(E)** Cortical activity during the lockout period for hit trials following a correct rejection (top) and hit trials not following a correct rejection (bottom) (n=39 sessions for both).

In our task, most target trials followed a correct rejection and shortened inter-trial interval (80%), while a minority of target trials was not preceded by a correct rejection and followed a long inter-trial interval (20%). It is possible that the mice in our task implemented a strategy of using the distractor stimulus to orient attention to the target whisker field rather than solely attending the target stimulus. To determine the likelihood of this strategy, we compared behavioral performance on target trials (Figure 7D) and hit-related neural activity (Figure 7E) separately according to the presence of a preceding correction rejection. The similar behavioral performance (hit rate, paired t-test, t(38)= −0.49, p=0.63; reaction time, paired t-test, t(38)=0.29, p=0.77) and neural activity suggest that the distractor stimulus was not utilized to enhance target detection.

Next, we sought to confirm our Ca^2+^ imaging findings with local field potential (LFP) recordings, which have much higher temporal resolution. We recorded LFP signals from layer 5 of S1, wMC and ALM, in target-aligned and distractor-aligned hemispheres (not simultaneously recorded). We compared target-evoked responses in target-aligned cortices (Figure 8 A-C) to distractor-evoked responses in distractor-aligned cortices (Figure 8 D-F). We find that early post-stimulus activity, likely reflecting the initial feedforward sensory sweep, is similar in target-aligned and distractor-aligned S1 and wMC (Figure 8 G, H; peak 1, occurring within 30 ms post-stimulus). Late activity does diverge in S1 and wMC between target and distractor recordings. In ALM, notably, the large post-stimulus activity in target recordings is nearly absent in distractor recordings. These LFP data support our Ca^2+^ imaging findings of minimal subcortical filtering of the sensory response followed by robust attenuation across cortex.

**Figure 8:**
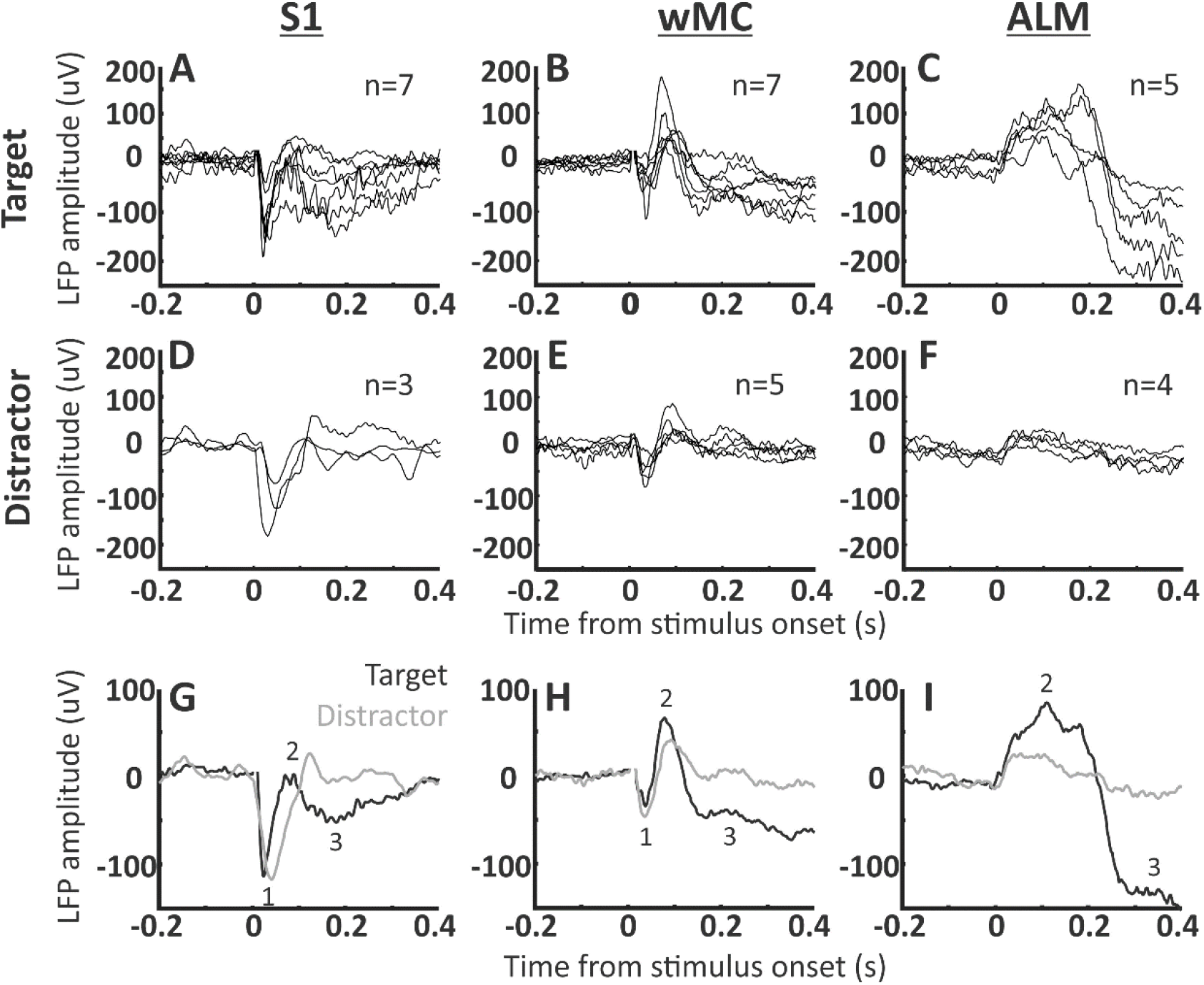
LFP signal transformation across S1, wMC and ALM. LFP signals were recorded from layer 5 of S1, wMC and ALM. (A-F) Each trace reflects average LFP signals for one session, across all target trials in target-aligned cortices (A-C) and across all distractor trials in distractor-aligned cortices (D-F). The count in each panel refers to the number of recorded sessions. (G-I) Target-aligned (black) and distractor-aligned (grey) LFP signals, averaged across sessions. We observed three distinct event-related potentials, two negative-going (1 and 3) and one positive-going (2). Event 1, which is large in S1, small in wMC and absent in ALM, likely reflects the initial feedforward sensory sweep. This event is similar in target and distractor recordings. Event 3, which is large in ALM and moderate in wMC and S1, is highly dissimilar between target and distractor recordings.

To further understand how neural activity relates to movements during the task, in four additional sessions we imaged whisker movements during task performance and analyzed task-aligned whisker motion energy. We present two example sessions in Figure 9, in which we plot whisker motion energy for target and distractor whiskers aligned to target and distractor stimulus trials. We find that whisker movements increase on target trials in both target and distractor whiskers approximately 100 ms after stimulus onset (Figure 9 A, B, E, F) (latency, n=4; target whiskers: 104 +/− 12 ms; distractor whiskers: 129 +/− 8 ms). This increase in bilateral whisker movements is before the onset of licking (>200 ms, due to lockout window), and therefore appears to be part of a reward motor sequence (Musall et al., 2019). Whisker motion energy on distractor trials remained at pre-stimulus levels, or increased late in the trial (Figure 9G), for target and distractor whiskers. In comparing the onset of neural signals (LFP) to the onset of behavior (whisking and licking) we find that the cortical signals precede overt behavior. Our data are therefore consistent with activation of wMC and ALM triggering a whisking and licking response sequence.

**Figure 9:**
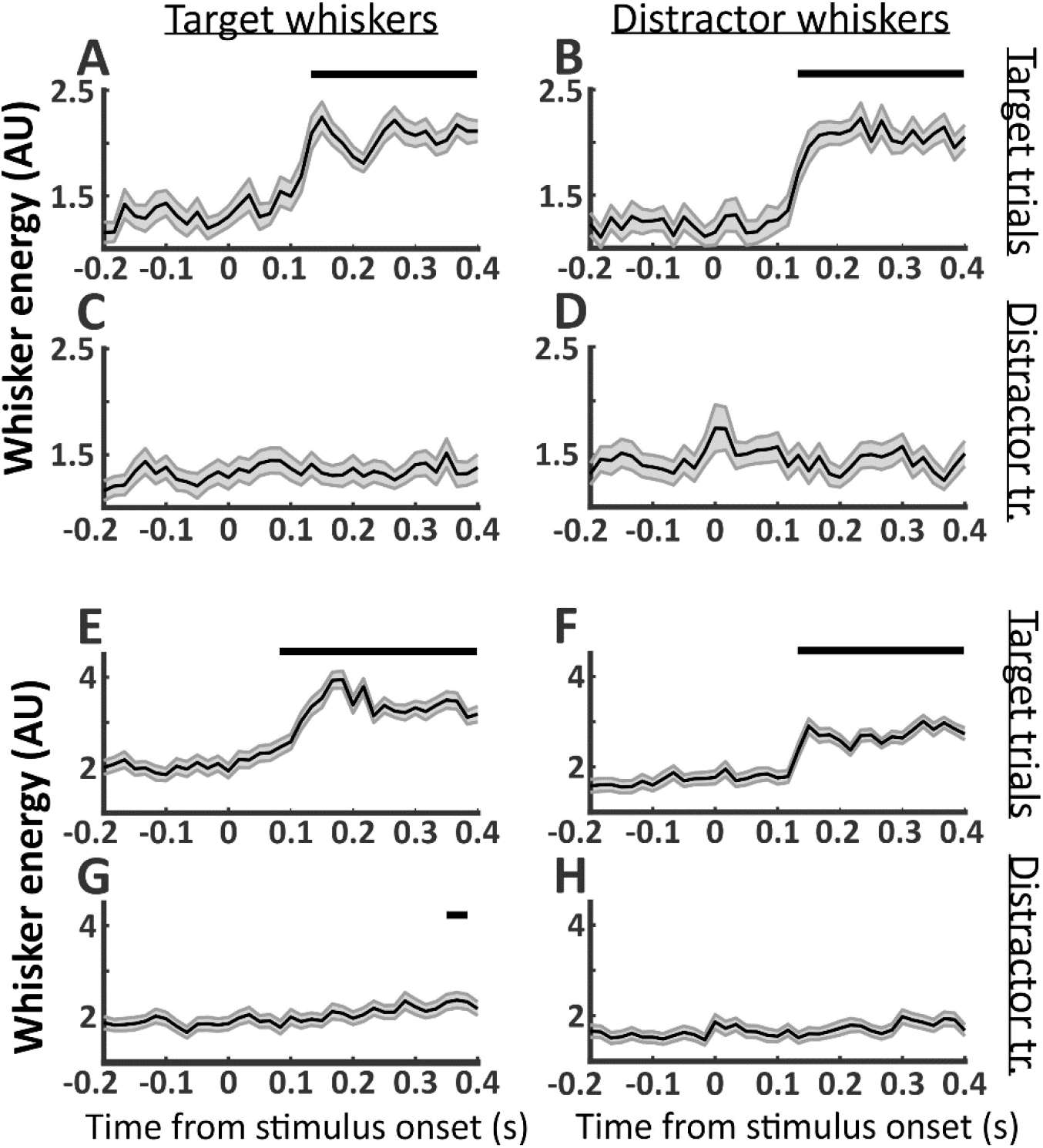
Bilateral whisker movements on target trials. Whisker energy was calculated over the target or distractor whisker fields. Significant changes from pre-stimulus whisker movements are indicated as black bars above the data. Two example sessions are shown, session 1 (A-D) and session 2 (E-H). (A, E) Target whisker energy on target trials; (B, F) distractor whisker energy on target trials; (C, G) target whisker energy on distractor trials; (D, H) distractor whisker energy on distractor trials. Significant increases in whisker movements occurred for both target and distractor whiskers approximately 0.1 seconds after target stimulus onset (A, B, E, F). Target and distractor whisker movements to distractor stimuli were either non-significant throughout the trial (C, D, H) or delayed (G).

We conducted additional analyses of the widefield imaging data to determine whether the observed propagation to frontal cortex for target stimuli is predictive of response initiation. The alternative hypothesis is that propagation reflects a learned stimulus association that may be independent of responding. To distinguish between these hypotheses, we calculated choice probability (Britten et al., 1996) for each pixel (Figure 10), comparing activity on hit trials (response present) versus miss trials (response absent). The average spatial map of target stimulus choice probability is shown in Figure 10A (left). This analysis revealed pixels with modest positive (increased on hit trials) and negative (increased on miss trials) values of choice probability. However, none of the pixel values were significantly different than zero after correction for multiple comparisons (Figure 10A, right).

**Figure 10:**
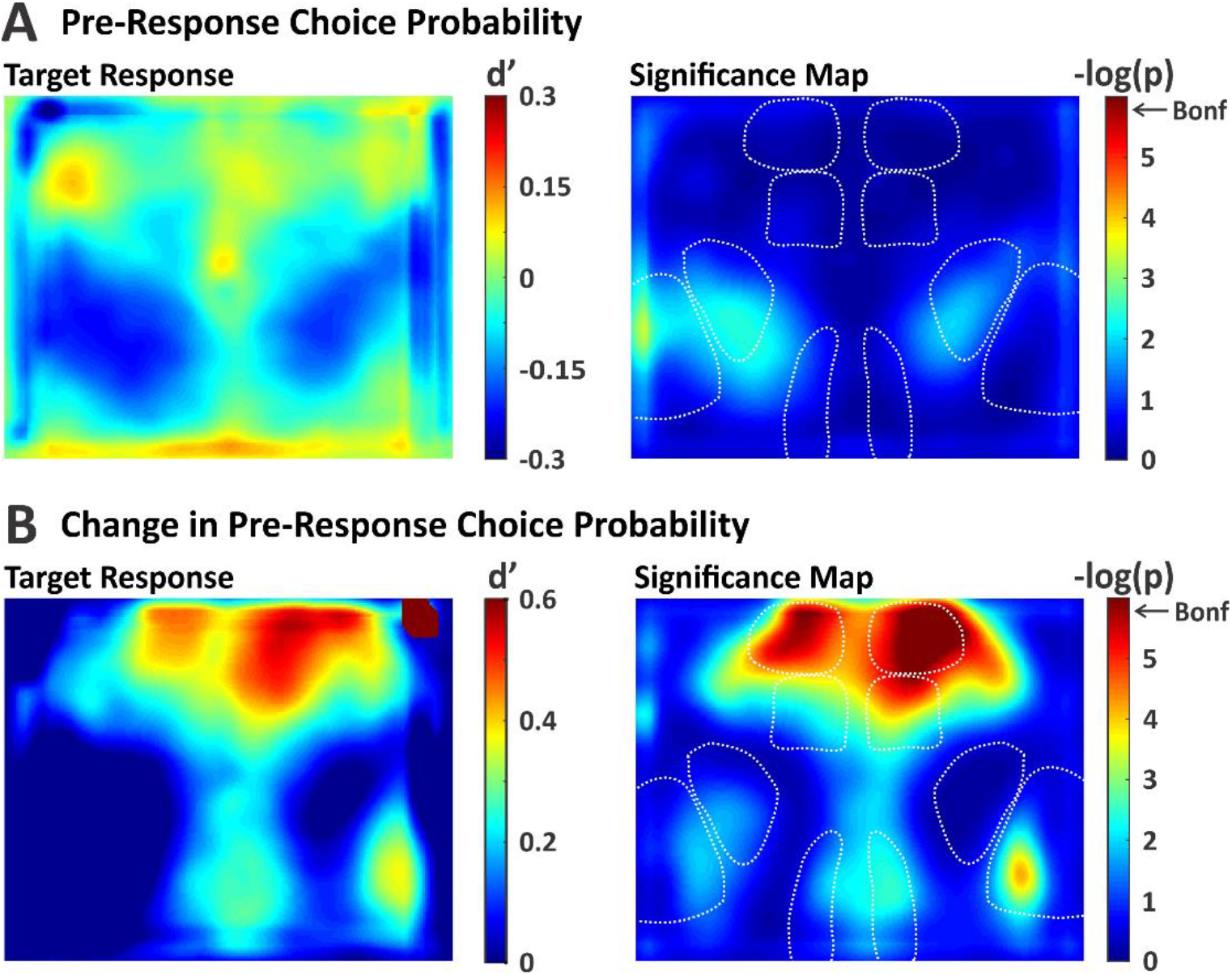
Spatial Maps of Choice Probability. We quantified choice probability as the separation between response absent and response present d’, computed pixel-by-pixel. **(A)** Choice probability map (left) and significance map (right) during the last frame of the lockout period. None of the pixels reached significance after correcting for multiple comparisons (Bonferonni). **(B)** Same as in [A], except with choice probability computed on the difference in activity between the two lockout frames. With this approach, significant choice probability was observed in target-aligned wMC and bilateral ALM.

We reasoned that hit versus miss outcomes may depend on both the state of the mouse as well as the strength of the stimulus-evoked responses. In order to isolate the latter component, we recalculated choice probability based on the difference in activity between early and late lockout period activity (see Methods). With this method, we observed large and significant choice probability values in target-aligned wMC and bilateral ALM (Figure 10B). We also observed focal increases in choice probability in target-aligned S1 and RSP, but these regions did not reach statistical significance after correction for multiple comparisons (Figure 10B). Thus, cortical activation of frontal cortex on target trials is predictive of response initiation.

### Quantification of functional connectivity across cortex

Finally, we sought to determine whether the differences in propagation for target versus distractor stimuli are reflected in the correlation patterns, or ‘functional connectivity’, between sensory and motor cortices. To do this, we created pixel-by-pixel correlation maps for S1, wMC, and ALM in target-aligned or distractor-aligned hemispheres as seed regions of interest (ROIs). We show the correlation maps for the full trial data, which includes the pre-stimulus, peri-stimulus and lockout, and response windows for all stimulus trial types (Figures 11A-F). The most striking findings are regional structure and symmetry. The spatial correlation patterns are highly similar for wMC and ALM seeds, which are quite different from S1 seeds (compare Figures 11A/D with 11B/E, C/F). This regional structure illustrates that the correlation values reflect local neural activity rather than global imaging artifacts. Regarding symmetry, for all three cortical regions, the target-aligned and distractor-aligned seed maps are qualitatively extremely similar (compare Figure 11A-C with 11D-F).

**Figure 11:**
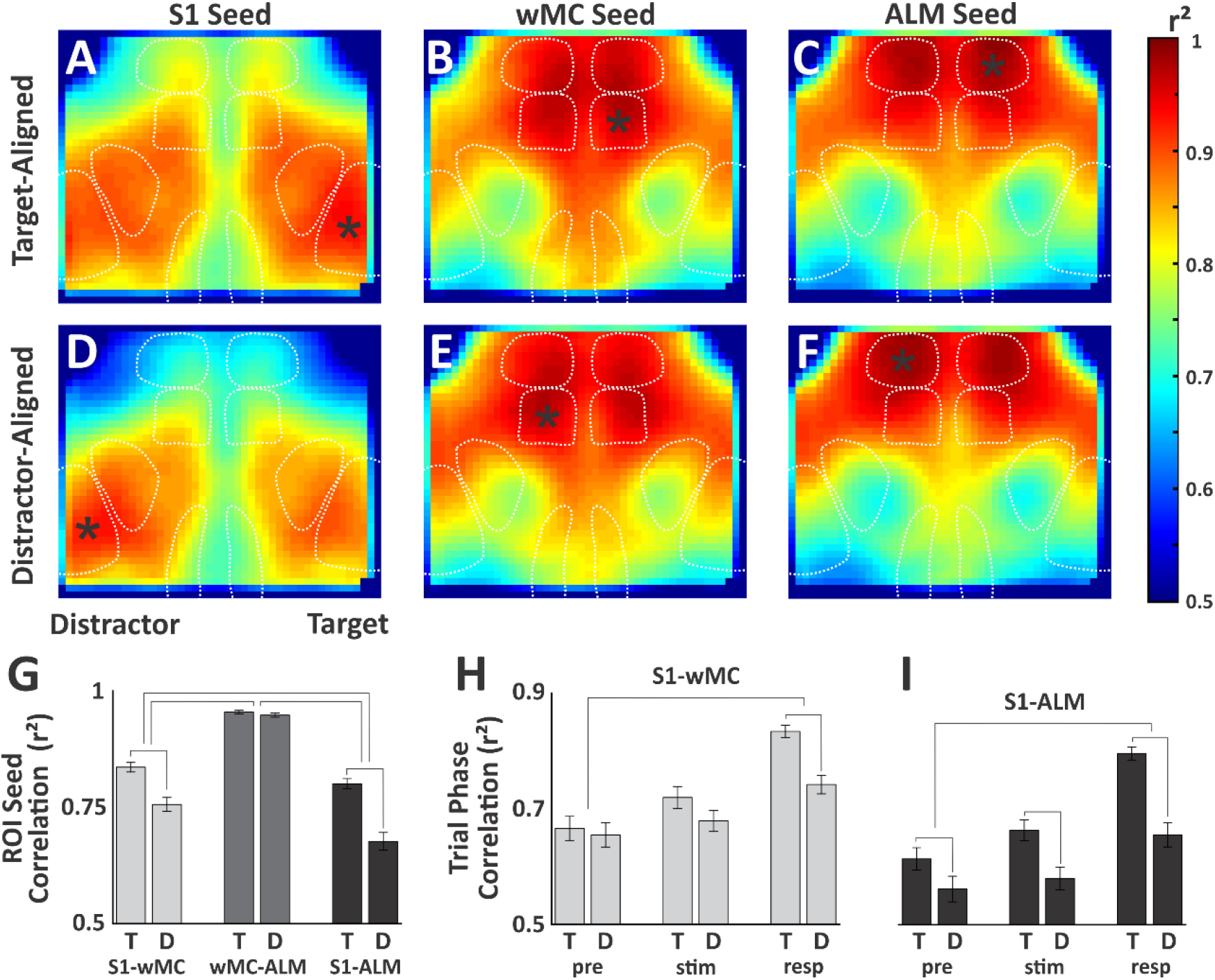
Spatial Correlation Analysis. **(A-F)** Correlation maps for full trial data. Seed regions of interest (marked by asterisk) included S1, wMC and ALM, in target-aligned **(A-C)** and distractor-aligned **(D-F)** cortices. **(G)** Summary data of average pairwise correlation values between S1 -wMC, wMC-ALM and S1-ALM. Statistical comparisons were made between target-aligned (T) and distractor-aligned (D) correlations, with statistical significance denoted by lines connecting adjacent columns. Statistical comparisons were also made based on the differences in target-aligned and distractor-aligned correlations between regions, with statistical significance denoted by lines connecting pairs of columns. (H, I) Similar structure as in [G], except for comparisons of target-aligned and distractor-aligned correlations (H, S1-wMC; I, S1-ALM) for different trial phases (pre, pre-stimulus; stim, stimulus and lockout; resp, response).

Despite these similarities, we do find significant differences in correlations between S1 and wMC (r^2^, target 0.84±0.01, distractor 0.76±0.01, paired t-test, t(38)=6.2, p=2.9e^-7^, n=39) and between S1 and ALM (r^2^, target 0.80±0.01, distractor 0.68±0.02, paired t-test, t(38)=8.1, p=8.6e^-10^, n=39). (Figure 11G). The largest differences in target-aligned and distractor-aligned correlations are between S1 and ALM (ANOVA, F(2,37)=30.3, p=1.6e^-8^, with post-hoc Tukey comparison) (Figure 11G). To determine whether these differences are persistent or related to specific phases of the task, we ran the correlation analyses separately for the pre-stimulus, peri-stimulus and lockout, and response windows. We found that differences in target-aligned versus distractor-aligned S1 to wMC (Figure 11H) and S1 to ALM (Figure 11I) correlations were significantly larger in the response phase compared to the pre-stimulus phase (ANOVA with post-hoc Tukey comparison, S1-wMC, p=0.0004; S1-ALM, p=0.0005). However, even in the pre-stimulus phase, there was a small yet significant increase in S1 to ALM correlation in target-aligned compared to distractor-aligned hemispheres (paired t-test, t(38)=3.4, p=0.0018) ((Figure 11I). Overall, these data are inconsistent with large, global changes in synaptic plasticity or functional connectivity driving task performance, but rather implicate more focal, possibly pathway-specific, adaptations.

## Discussion

We developed a Go/NoGo selective detection task to study the neural processes of sensory selection in the mouse somatosensory whisker system. Mice learned to respond to target whisker deflections and ignore contralateral, distractor whisker deflections, achieving expert performance within 2 to 3 weeks of training (Figure 2). The main finding of this study is robust attenuation of distractor compared to target stimulus processing between mono-synaptically coupled cortical regions S1 and wMC (Figures 3–6). We interpret this observation as reflecting the presence of an intra-cortical attenuating filter, suppressing higher order processing of unattended stimuli (Treisman, 1964).

We note important differences between our study and previous studies of the neural correlates of sensory selection. In our task, target and distractor receptive fields were assigned at the onset of training and remained constant throughout the learning process. This contrasts with previous studies in primates, in which target and distractor assignments are cued each block or trial. Moreover, our target and distractor stimuli were always across hemispheres, rather than varying in proximity. Yet, despite differences in training, species, sensory modality, stimulus details, and recording technique, we do note remarkable similarities with previous studies. As in the primate visual system, we observe progressive distractor suppression along the cortical hierarchy (Figure 6D) (Moran and Desimone, 1985; Tootell et al., 1998; Treue, 2001). Comparing modulation amplitudes between studies is problematic, because they vary widely depending on task and stimulus details. However, generally, within thalamus and primary visual cortex, attentional modulations of approximately 10% have been reported (Motter, 1993; Tootell et al., 1998; McAlonan et al., 2008), which is similar to the 8.8% average modulation we observe in primary somatosensory cortex. Within higher order sensory cortices, attentional modulations of 50 to 65% have been reported (Moran and Desimone, 1985; Tootell et al., 1998), which is similar to the 61.0% and 72.1% average modulations we observe in wMC and ALM, respectively.

What is the nature of distractor suppression? One possibility is that suppression is reactive, that once a distractor is detected, another brain region initiates an inhibitory brake to prevent further processing. This type of transient activation is observed, for example, in prefrontal cortex during stop-signal reaction time tasks at the detection of a ‘stop’ signal (Hanes et al., 1998; Aron and Poldrack, 2006). A second possibility is that suppression is proactive, already deployed in the initial conditions of the brain regions receiving the distractor stimulus. Insofar as we do not observe additional transient activations for distractor stimuli, our data support the second explanation of proactive suppression.

Given our localization of an attenuating filter between S1 and wMC, there are multiple possible mechanisms for implementing this filter. The most direct mechanism would be bidirectional modulation of the S1-wMC intra-cortical projection pathway. Previous studies of whisker detection have identified increased sensory processing with learning in wMC and in specific S1-wMC projection neurons (Chen et al., 2015; Le Merre et al., 2018). Whether this pathway decreases in strength when aligned with a distractor has not been studied. However, such a finding of bidirectional modulation would provide strong evidence for involvement of this pathway in specific stimulus selection, rather than general task engagement. Additionally, regulated propagation between S1 and wMC may involve subcortical loops through the striatum (Alloway et al., 2006) or posterior medial thalamus (Kleinfeld et al., 1999), or cortical feedback projections from PFC to wMC or from wMC to S1 (Xu et al., 2012; Zagha et al., 2013). For example, wMC to S1 feedback may strengthen (target-aligned) or weaken (distractor-aligned) the reciprocal S1 to wMC feedforward pathway. Strengthening or weakening may occur through feedback targeting of excitatory, inhibitory or disinhibitory S1 neurons (Rocco and Brumberg, 2007; Petreanu et al., 2009; Lee et al., 2013; Zagha et al., 2013; Kinnischtzke et al., 2014). Our task provides an excellent platform for studying the plasticity and cellular/circuit contributions of each of these mechanisms towards target enhancement and/or distractor suppression. Alternatively, our findings are inconsistent with strong reductions in ascending sensory drives to distractor-aligned S1 (Figures 3–6 and 8) or large, global reductions in the structural or functional connectivity between this region and the rest of cortex (Figure 11).

While our study identifies a sensory filtering process distal to S1, other studies have identified sensory gating in S1 and earlier subcortical structures. Previous studies of the rodent whisker system have examined differences in sensory processing during periods of whisking versus non-whisking. In general, these studies find reductions in sensory responses during whisking (Fanselow and Nicolelis, 1999; Crochet and Petersen, 2006; Ferezou et al., 2007; Lee et al., 2008; Chakrabarti and Schwarz, 2018), which is already present in the first sensory brainstem relay (Chakrabarti and Schwarz, 2018). This sensory gating process is likely mediated by both top-down cortical (Lee et al., 2008; Chakrabarti and Schwarz, 2018) and neuromodulatory (Eggermann et al., 2014) inputs. Thus, modulations of sensory processing may occur all along the ascending sensory pathway, including brainstem, thalamus and cortex. Why different behavioral contexts engage different mechanisms of sensory gating is currently unknown.

Finally, we currently do not know how wMC contributes to the sensory selection process. This cortical region has been studied extensively with respect to whisking, specifically in establishing its set-point, initiation, and amplitude modulation (Carvell et al., 1996; Hill et al., 2011). Consistent with this, we find that wMC activation on target trials correlates with bilateral increases in whisking (Figure 9). Alternatively, more recent studies have demonstrated roles for this same region in orienting behaviors and action suppression (Erlich et al., 2011; Zagha et al., 2015; Ebbesen et al., 2017). Our study demonstrates, at the representational level, a possible additional function of regulating the propagation of sensory processing for sensory selection. And yet, wMC is only one of the many routes by which a whisker stimulus can initiate a motor output (Kleinfeld et al., 1999). Defining how wMC contributes to sensory-motor processing in this task and in other behavioral contexts will be a major focus of future investigations.

## Acknowledgements

This work was supported by the Whitehall Foundation (Research Grant 2017-05-71 to E.Z.) and the National Institutes of Health (R01NS107599 to E.Z.). We thank Trevor Stavropoulos for assistance with Arduino programming. We thank Shaida Abachi for collecting whisker imaging data. We thank Hongdian Yang, Martin Riccomagno, Aaron Seitz and Megan Peters for many helpful discussions throughout the project. The authors declare no competing financial interests.

